## Supplemental Information for "Higher-order epistasis shapes natural variation in germ stem cell niche activity"

### SUPPORTING INFORMATION

**This PDF file includes:**
Legend for Data S1 (separate file)
Figs. S1 – S4
Tables S1 – S14

**Legend for Data S1. (separate .xlsx file)**
**Each of the items listd is a separate worksheet in Data S1.xlsx**

**Fig1C:** Data used to determine the relationship between progenitor zone size (area in microns) and progenitor cell number.

**Fig1D**: Young adult (1-10 eggs *in utero*) progenitor zone size among select wild isolates.

**Fig2A**: Progenitor zone cell number in JU1200 and JU751 at mid L4 (mL4), young adult (YA), 24 hours post mid L4 (mL4+24), and 24 hours post young adult (YA+24).

**Fig2B**: Number of EdU + cells in the progenitor zone of JU1200 and JU751 at mid L4 (mL4).

**Fig3A_corr_factor**: Raw QTL mapping data—progenitor zone area in pixels—was corrected for block effects in preparation for mapping. Correction factors (Corr_factor) were derived from the least square means of the model in Table S5. Raw progenitor zone area was divided by the correction factor for each block to produce the corrected data (Corr_PZArea) in pixels. QTL mapping was performed with the Corr_PZ data. Values were scaled to square microns (in um^2) for the purpose of presentation in Figure 3A.

**Fig3A_mapping**: Data frame used as input for QTL mapping in R/qtl.

**Fig4C**: Young adult (1-10 eggs *in utero*) progenitor zone size among JU1200 NILs

**Fig4D**: Young adult (1-10 eggs *in utero*) progenitor zone size among JU751 NILs

**Fig5B**: Young adult (1-10 eggs *in utero*) progenitor zone size among parental (JU1200, JU751) and CRISPR derived reciprocal allelic replacement lines containing *lag-2p(cgb1008)* and lag-2(cgb1007) in the non-native backgrounds. Progenitor zone area was measured in pixels (Area_pxl) and scaled to area in square microns (PZSize_um) for graphical presentation.

**Fig5D-F**: Number of *lag-2* smFISH puncta counted in a distal tip cell at mid L2 (mL2), early L3 (eL3), and mid L4 (mL4) in JU1200 and JU751.

**Fig6**: Young adult (1-10 eggs *in utero*) progenitor zone size among parental (JU1200, JU751) lines, CRISPR derived reciprocal allelic replacement lines, and near isogenic lines. Progenitor zone area was measured in pixels (Area_pxl) and scaled to area in square microns (PZSize_um) for graphical presentation. Includes the genotypes of the genetic background, central portion of chromosome II QTL (C2), and *lag-2p* deletion (C5).

**FigS4:** Number of *lag-2* smFISH puncta counted in a distal tip cell at young adult (YA, 1-10 eggs *in utero*) in JU1200 and JU751.

**chrIIQTL variants**: A table listing all known SNPs and short variants between JU1200 and JU751 within the reduced 2.04 Mb chromosome II QTL interval. Data was collected from CeNDR.
**chrVQTL variants**: A table listing all known SNPs and short variants between JU1200 and JU751 within the 2.4 Mb chromosome V QTL interval. Data was collected from CeNDR.

**INDEL variants in QTL**: A table listing large INDEL variants between JU1200 and JU751 in the reduced 2.04 Mb chromosome II QTL interval and the 2.4 Mb chromosome V QTL interval. Data was collected from CeNDR.

### Supporting Figures

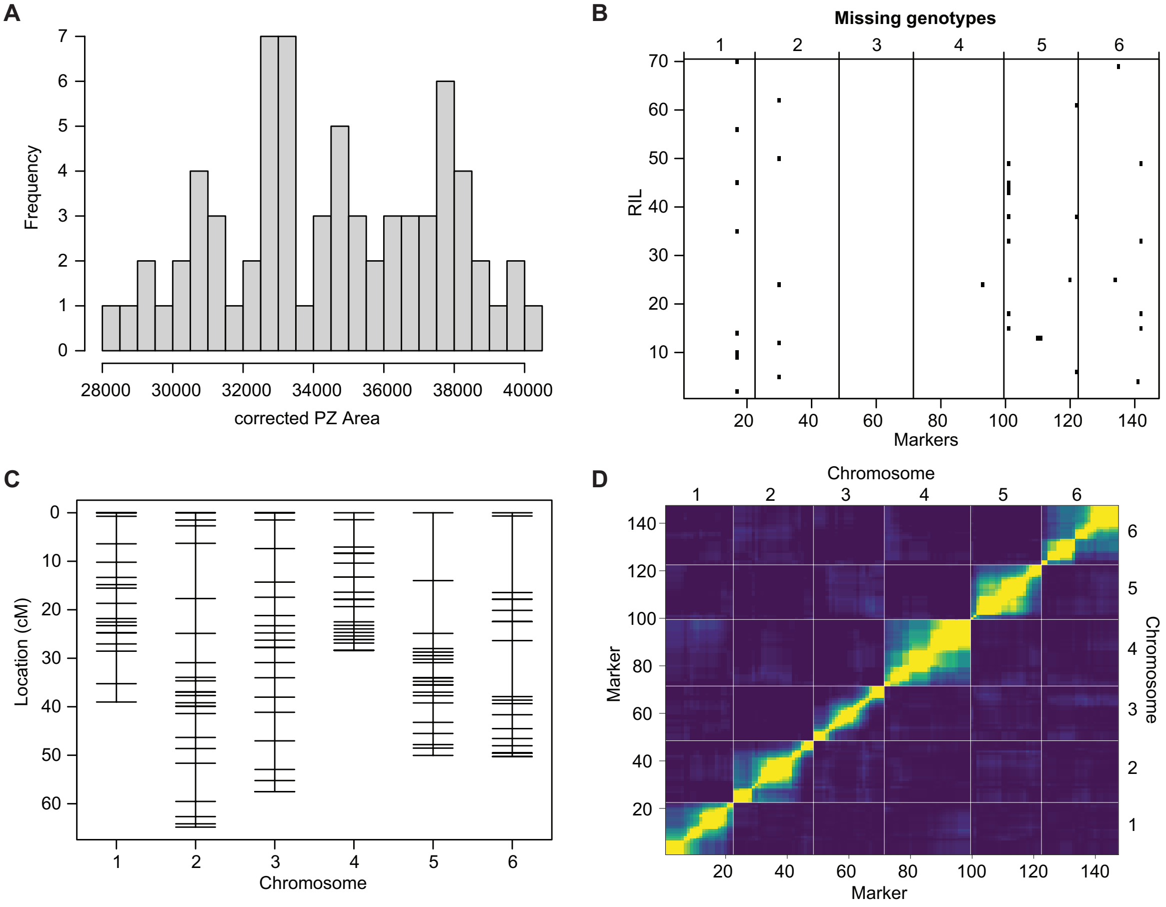


**Figure S1. Additional data from QTL mapping of RILs.**
A) Phenotype distribution for all RILs. Corrected means for each RIL were calculated using correction factors derived from the least square means of a data set containing only the parental lines from each block (S3-S8). See Experimental Procedures and Table S5 for more detail. B) Missing genotypes for each RIL across all markers. C) Genetic markers cover the entire genome. D) Heat map of the recombination fractions (top) and LOD scores (bottom) for the genetic map, yellow corresponds to less recombination between markers while purple corresponds to more recombination between markers. LOD scores are derived by testing whether the recombination fraction equals 0.5, as per the est.rf() function in the R/qtl package.

JU751cV del3.2 fwd1 / JU751cV del3.2 fwd2

TCCGTTATGACTCTATTGCTCCTAATCCCCCTTTTGTTTCCGGTTCGGCCTGAAACAAGAATAACAACAGGTGACTCGATA

ACTGTTAGGAAAATAAATTTTTTGGTCCCAATATTTCGGTATCCATTTCAAAACATCTTTTAAAATTGTCAAGACAGACAATT

JU751cV del3.2 F3

ACTTCTGCTTGAAATGAAGGACACATGTCCCCTCAATTCCTCATAAAACAAATAGAGGGGTGTCGAAATGATAAATGTTGC

G->A in repair

AAGTGAGGTGCAGATGGCAATGAAATTTATATATGTTT**CTTTCATTCCATTCTTGTGGGGCAATTTCAAACGTTTGATAAC**

HLH-2 site / JU751cV del3-2 F5/R5

**TCACAGTGGTCTAGATTTGCAGAACATCTGTGTTAGGTCAAGCAGAGCGAGAATTTGAGCATTTGAACTTTAGGGTCC**

G->A in repair / G->T in repair

**TGGGGGGGGAGGGGGGGGGGGGGGG**GGGGTGTCTAGACAGTCAGCGGCCCATAAGGGTTCATATGGTTTGACGG

TGCATTGAACTATTTTTTAAGGAAAAAATTGAGATAAAGGCCATTTTTCCAAGAAAGCATCTAGCTTTTTCAACTGACACTA

JU751cV del3-2 R4

CAAATTTCCCTATAACTTTCTGAAATTATCAGTGAAAAGTTGCCCAAATTTGAGACTTTAGCTAAAAATAGACAATTTTTTC

SNP C->A

CAAAACTTTGAGCGGCCATAACTTTTTTTTGAAAAACTTTCGGAACGTCTCATTACAAAATTTGGTTGCTTTGAGCCATTTT

GAATCCGAAAAAGTCTCAAAAAGCAAAGTCTTAAATTTCGGTACTCCACCTTTAAAAAAAATAGGGTTTCCGCCGAAATTT

CTGGGTTCCAATAATATCTGTTACCATTTCCTTGCTCTGCGACGAGCAGCCACCAGACAAAAATCTCCTAATACCCCTCTT

JU751cV del3.2 rev1

CATGCCTTTGTTCTTCCGGTTTGGCGGTCTGAAAAAGAGCACCCCTCCCGGCGGGGGGCTCTCGGTATCTGTTCCGCAA

JU751cV del3.2 rev2

AAACCGTCATATGGAGGGCTATCGGGAGACGGGAAGAGTTGGAAATACGGGAAATGGTGATG

**Figure S2. Portion of *lag-2* promoter sequence in JU1200 containing the 148 bp deleted in JU751.** The deletion is bolded and underlined. The HLH-2 binding site, genotyping primers, CRISPR induced mutations, and SNPs are annotated. See Table S13 for primer sequences.

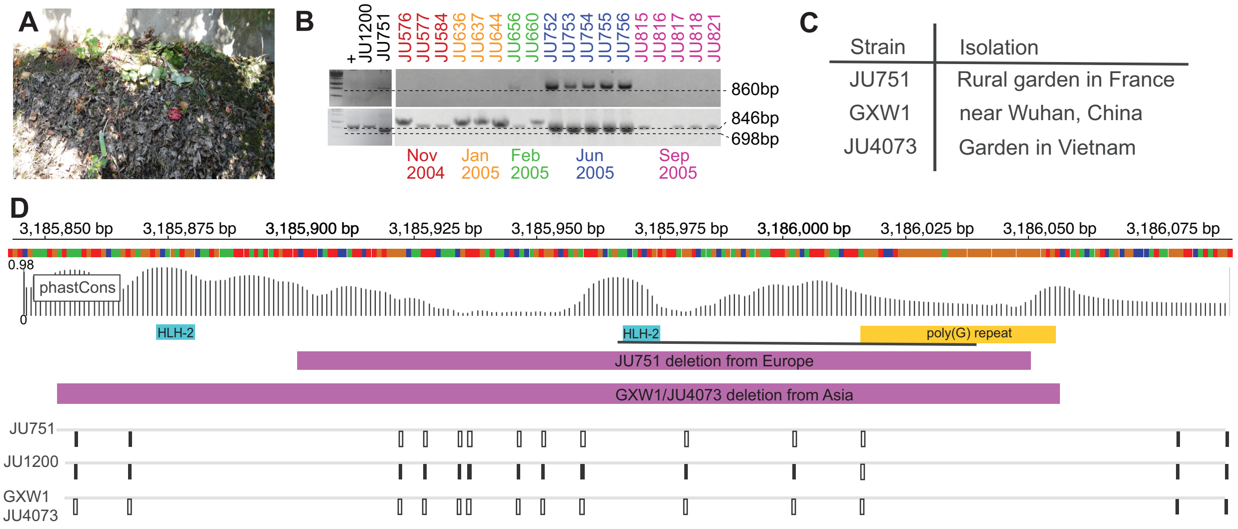


**Figure S3. The JU751 deletion arose in the wild and is a recent, unique event.** A) A photograph of the compost pile in Le Perreux-sur-Marne, France from which JU751 and the strains in panel B were isolated. B) Strains isolated from the compost pile in panel A during the months shown were genotyped for the presence of the *lag-2(cgb1007)* deletion. PCR results from two sets of primers spanning the 148 bp deletion are shown. Single PCR across the long polyG repeat is not successful leading to lack of a band in JU1200 and a band of 860 bp in JU751 (top gel). Strains JU752-756 isolated at the same time and place also give an 860 bp band while other strains isolated from the same place at different times of the year give no band. A nested PCR reaction generates an 846 bp band in JU1200 and a 698 bp band in JU751. JU752-756 give the 698 bp band while all others give an 846 bp band or larger. C) Isolation locations for JU751 and strains containing a larger, but similar deletion (Cook et al. 2017). D) Variant annotation data from CeNDR showing the deletion region in JU1200, JU751, GXW1, and JU4073. The extent of the deletions are marked by the magenta bars. The HLH-2 binding sites are marked by cyan bars. The polyG repeat is marked by a gold bar. Variants from ~1500 annotated *C. elegans* strains are shown as dark gray solid bars. "NO-CALL" variants are show as empty bars. Further confirmation that the deletion is only present in JU751 can be found in the CeNDR database (https://elegansvariation.org). Reference sequence data is from genome WS276.

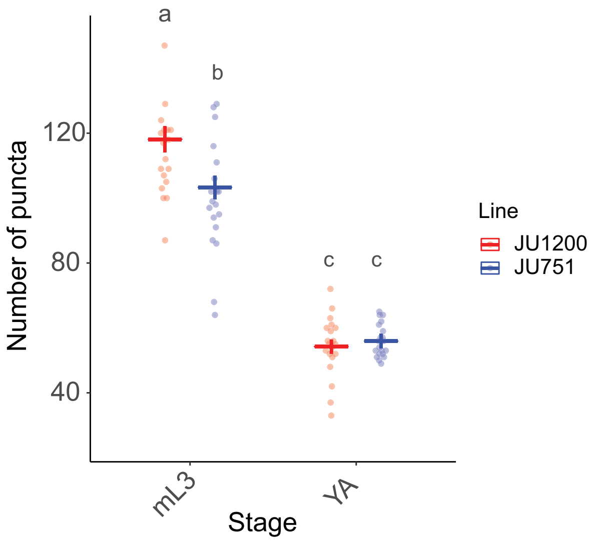


**Figure S4. *lag-2* expression in the DTC of JU1200 and JU751 at midL4 and young adult.**
Data from quantification of *lag-2* transcripts via smFISH at midL3 and young adult. Crossbars and error bars represent estimated marginal means +/- standard error from a generalized linear model with a negative binomial distribution. Different bold lowercase letters indicate groups that are significantly different (p<0.05). For statistical models and results see Table S12.

### Supporting Tables

**Table S1 (accompanies Figure 1C). Statistical testing for correlation between the number of progenitor zone nuclei (“Cells”) and progenitor zone size (“PZSize”).** Data from multiple strains and two adult stages (L4+24h, YA+24h) were fitted to a gaussian linear model, PZSize (μm$^2$) ~ Cells+Strain+Stage. For data representation (strain and stage aggregated, as they are not significant in the model below), see Figure 1C.

**A) Correlation between PZ cell number and PZ size: Gaussian linear model**

Residual standard error: 321.8 on 83 degrees of freedom
Multiple R-squared 0.7662
**Adjusted R-squared 0.758**
F-statistic 90.69 on 3 and 83 degrees of freedom
**p-value <2.2E-16**

| *Source* | *Estimate* | *SE* | *t-value* | *Pr>\|z\|)* | |
|---|---|---|---|---|---|
| *Intercept* | 414.842 | 222.83 | 1.862 | 0.0662 | |
| *Cells* | 12.2113 | 0.9135 | 13.367 | <2.2E-16 | *** |
| *StrainJU751* | 115.667 | 104.17 | 1.110 | 0.2700 | |
| *StageYA+24* | 54.7432 | 82.777 | 0.661 | 0.5102 | |

**Table S2 (accompanies Figure 1D). Statistical testing for differences in the size of the progenitor zones among selected wild isolates of *C. elegans* and *C. briggsae.*** A) Raw data (PZ area in pixels) were fitted to a generalized linear mixed model, PZArea ~ Strain + (1|Exp), with a negative binomial distribution (log link) and a dispersion parameter of 37.5. The R software package, ‘emmeans’ was used to obtain estimated marginal means on the response scale (B) and pairwise contrasts (C) with Tukey corrected p-values. Area in pixels was scaled to μm$^2$/1000 for representation in Figure 1D.

**A) PZ Size in wild isolates: Negative binomial model**

| *Source* | *Estimate* | *SE* | *Z-value* | *Pr>\|z\|)* | |
|---|---|---|---|---|---|
| *Intercept* | 10.41613 | 0.0272 | 382.9 | < 2e-16 | *** |
| *CB4856* | -0.14976 | 0.03268 | -4.6 | 4.59E-06 | *** |
| *CX11262* | -0.04457 | 0.03417 | -1.3 | 0.192034 | |
| *HK104* | -0.10483 | 0.03268 | -3.2 | 0.001337 | ** |
| *JU1200* | 0.25508 | 0.02981 | 8.6 | < 2e-16 | *** |
| *JU1491* | 0.15314 | 0.03425 | 4.5 | 7.77E-06 | *** |
| *JU751* | 0.06414 | 0.03001 | 2.1 | 0.032583 | * |
| *JU775* | 0.08888 | 0.03411 | 2.6 | 0.009177 | ** |
| *MY16* | 0.03845 | 0.03419 | 1.1 | 0.260849 | |
| *N2* | 0.10019 | 0.02993 | 3.3 | 0.000817 | *** |

**B) PZ Size in wild isolates: Emmeans**

| *Strain* | *response* | *SE* | *df* |
|---|---|---|---|
| *AF16* | 33393.7851 | 908.415431 | 1139 |
| *CB4856* | 28749.1438 | 782.118305 | 1139 |
| *CX11262* | 31938.0063 | 659.990888 | 1139 |
| *HK104* | 30070.3221 | 818.044304 | 1139 |
| *JU1200* | 43096.8579 | 707.047176 | 1139 |
| *JU1491* | 38919.9543 | 809.491218 | 1139 |
| *JU751* | 35605.6964 | 595.209483 | 1139 |
| *JU775* | 36497.6877 | 751.01212 | 1139 |
| *MY16* | 34702.6497 | 718.646237 | 1139 |
| *N2* | 36912.7142 | 614.851536 | 1139 |

**C) PZ Size in wild isolates: Contrasts**

| *Contrast* | *ratio* | *SE* | *df* | *t-ratio* | *p -value* | |
|---|---|---|---|---|---|---|
| *AF16 / CB4856* | 1.16155755 | 0.03795836 | 1139 | 4.58283737 | 0.00021846 | *** |
| *AF16 / CX11262* | 1.04558139 | 0.03572386 | 1139 | 1.30458463 | 0.95269564 | |
| *AF16 / HK104* | 1.11052303 | 0.0362901 | 1139 | 3.20796461 | 0.04458804 | * |
| *AF16 / JU1200* | 0.77485429 | 0.02310083 | 1139 | -8.5559722 | 1.27E-13 | *** |
| *AF16 / JU1491* | 0.85801193 | 0.02938501 | 1139 | -4.4714502 | 0.00036324 | *** |
| *AF16 / JU751* | 0.9378776 | 0.02814523 | 1139 | -2.1371846 | 0.50138738 | |
| *AF16 / JU775* | 0.91495619 | 0.03121234 | 1139 | -2.6053949 | 0.21612601 | |
| *AF16 / MY16* | 0.96228344 | 0.03290344 | 1139 | -1.1243861 | 0.98222963 | |
| *AF16 / N2* | 0.90466892 | 0.02708016 | 1139 | -3.3469298 | 0.0288436 | * |
| *CB4856 / CX11262* | 0.90015462 | 0.03075644 | 1139 | -3.0785783 | 0.06535972 | |

| | | | | | | |
|---|---|---|---|---|---|---|
| *CB4856 / HK104* | 0.95606371 | 0.03124402 | 1139 | -1.3748753 | 0.93477085 | |
| *CB4856 / JU1200* | 0.66708213 | 0.01988891 | 1139 | -13.578571 | 3.41E-14 | *** |
| *CB4856 / JU1491* | 0.73867363 | 0.02529901 | 1139 | -8.8439648 | 1.20E-13 | *** |
| *CB4856 / JU751* | 0.80743102 | 0.02423192 | 1139 | -7.127276 | 8.17E-11 | *** |
| *CB4856 / JU775* | 0.78769768 | 0.02687226 | 1139 | -6.9952016 | 2.03E-10 | *** |
| *CB4856 / MY16* | 0.82844233 | 0.02832821 | 1139 | -5.5040371 | 2.04E-06 | *** |
| *CB4856 / N2* | 0.77884123 | 0.02331494 | 1139 | -8.3495758 | 1.48E-13 | *** |
| *CX11262 / HK104* | 1.06211055 | 0.03628968 | 1139 | 1.76360516 | 0.75804118 | |
| *CX11262 / JU1200* | 0.74107505 | 0.01489957 | 1139 | -14.904163 | 3.41E-14 | *** |
| *CX11262 / JU1491* | 0.8206075 | 0.0173465 | 1139 | -9.3530432 | 1.29E-13 | *** |
| *CX11262 / JU751* | 0.89699148 | 0.01823029 | 1139 | -5.3488432 | 4.75E-06 | *** |
| *CX11262 / JU775* | 0.87506931 | 0.01830673 | 1139 | -6.3790696 | 1.16E-08 | *** |
| *CX11262 / MY16* | 0.92033336 | 0.01937291 | 1139 | -3.9439331 | 0.00338054 | ** |
| *CX11262 / N2* | 0.8652305 | 0.01759653 | 1139 | -7.1178928 | 8.72E-11 | *** |
| *HK104 / JU1200* | 0.69773815 | 0.02080256 | 1139 | -12.07178 | 3.41E-14 | *** |
| *HK104 / JU1491* | 0.77261966 | 0.0264613 | 1139 | -7.5321861 | 4.66E-12 | *** |
| *HK104 / JU751* | 0.84453683 | 0.02534509 | 1139 | -5.6302347 | 1.01E-06 | *** |
| *HK104 / JU775* | 0.82389663 | 0.02810683 | 1139 | -5.6782355 | 7.71E-07 | *** |
| *HK104 / MY16* | 0.86651372 | 0.02962966 | 1139 | -4.1901175 | 0.00123886 | ** |
| *HK104 / N2* | 0.81463319 | 0.02438598 | 1139 | -6.8487686 | 5.47E-10 | *** |
| *JU1200 / JU1491* | 1.10732036 | 0.02241568 | 1139 | 5.03591895 | 2.43E-05 | *** |
| *JU1200 / JU751* | 1.21039222 | 0.02118117 | 1139 | 10.9114687 | 1.09E-13 | *** |
| *JU1200 / JU775* | 1.18081064 | 0.02363407 | 1139 | 8.30378008 | 1.56E-13 | *** |
| *JU1200 / MY16* | 1.24188955 | 0.0250247 | 1139 | 10.7508008 | 1.19E-13 | *** |
| *JU1200 / N2* | 1.16753425 | 0.02039989 | 1139 | 8.86495347 | 1.19E-13 | *** |
| *JU1491 / JU751* | 1.09308224 | 0.02236466 | 1139 | 4.3499841 | 0.00062327 | *** |
| *JU1491 / JU775* | 1.06636767 | 0.02245004 | 1139 | 3.0522364 | 0.07045444 | |
| *JU1491 / MY16* | 1.12152688 | 0.0237557 | 1139 | 5.41466278 | 3.33E-06 | *** |
| *JU1491 / N2* | 1.05437802 | 0.02158697 | 1139 | 2.58630107 | 0.22521825 | |
| *JU751 / JU775* | 0.97556033 | 0.01974004 | 1139 | -1.2228218 | 0.96874758 | |
| *JU751 / MY16* | 1.02602242 | 0.02089853 | 1139 | 1.26124197 | 0.96180811 | |
| *JU751 / N2* | 0.96459166 | 0.01711941 | 1139 | -2.0312575 | 0.57694113 | |
| *JU775 / MY16* | 1.05172625 | 0.0220481 | 1139 | 2.40572088 | 0.32320127 | |
| *JU775 / N2* | 0.98875654 | 0.02002054 | 1139 | -0.5584271 | 0.99992588 | |
| *MY16 / N2* | 0.94012728 | 0.0191617 | 1139 | -3.0291396 | 0.07518765 | |

**Table S3 (accompanies Figure 2A). Statistical testing for differences in the number of progenitor zone nuclei between JU1200 and JU751 at L4 and adult stages.** Data were fitted to a generalized linear model with negative binomial distribution: Cells ~ Strain*Stage, with a dispersion formula ~ Strain*Stage, and log link (A, B). The R software package, 'emmeans' was used to obtain estimated marginal means on the response scale (C) and pairwise contrasts (D) with Tukey corrected p-values. For data representation, see Figure 2A.

**A) PZ cells in JU1200 and JU751: Negative binomial conditional model**

| *Source* | *Estimate* | *SE* | *Z-value* | *Pr>\|z\|)* | |
|---|---|---|---|---|---|
| *Intercept* | 5.213 | 0.0585 | 89.11 | <2e-16 | *** |
| *StrainJU751* | -0.038 | 0.0866 | -0.44 | 0.6573 | |
| *StagemL4+24* | 0.3053 | 0.0661 | 4.62 | 3.85e-6 | *** |
| *StageYA* | 0.4014 | 0.0637 | 6.30 | 2.9e-10 | *** |
| *StagemYA+24* | 0.0737 | 0.0855 | 0.87 | 0.3821 | |
| *StrainJU751:StagemL4+24* | -0.2628 | 0.1170 | -2.25 | 0.0248 | * |
| *StrainJU751:StageYA* | -0.0850 | 0.0945 | -0.90 | 0.3686 | |
| *StrainJU751:StageYA+24* | 0.2221 | 0.1294 | -1.72 | 0.0862 | |

**B) PZ cells in JU1200 and JU751: Negative binomial dispersion model**

| *Source* | *Estimate* | *SE* | *Z-value* | *Pr>\|z\|)* | |
|---|---|---|---|---|---|
| *Intercept* | 2.7645 | 0.3425 | 8.072 | 6.89e-16 | *** |
| *StrainJU751* | -0.2384 | 0.4751 | -0.502 | 0.6159 | |
| *StagemL4+24* | 1.6540 | 0.5717 | 2.893 | 0.0038 | ** |
| *StageYA* | 1.9473 | 0.5614 | 3.468 | 0.0005 | *** |
| *StagemYA+24* | -0.0864 | 0.4859 | -0.178 | 0.8589 | |
| *StrainJU751:StagemL4+24* | -1.7624 | 0.7473 | -2.358 | 0.0184 | * |
| *StrainJU751:StageYA* | -0.0974 | 0.7689 | -0.127 | 0.8992 | |
| *StrainJU751:StageYA+24* | -0.1461 | 0.6779 | -0.215 | 0.8294 | |

**C) PZ cells in JU1200 and JU751: Emmeans**

| *Strain* | *Stage* | *response* | *SE* | *df* |
|---|---|---|---|---|
| *JU1200* | *mL4* | 183.686 | 10.427 | 140 |
| *JU751* | *mL4* | 176.762 | 11.287 | 140 |
| *JU1200* | *mL4+24* | 249.263 | 7.662 | 140 |
| *JU751* | *mL4+24* | 184.441 | 13.367 | 140 |
| *JU1200* | *YA* | 274.398 | 6.896 | 140 |
| *JU751* | *YA* | 242.551 | 6.840 | 140 |
| *JU1200* | *YA+24* | 197.932 | 12.331 | 140 |
| *JU751* | *YA+24* | 152.540 | 11.181 | 140 |

**D) PZ cells in JU1200 and JU751: Contrasts**

| *Contrast* | *ratio* | *SE* | *df* | *t-ratio* | *p -value* | |
|---|---|---|---|---|---|---|
| *JU1200 mL4 / JU751 mL4* | 1.039 | 0.090 | 140 | 0.4437 | 9.998E-01 | |
| *JU1200 mL4 / JU1200 mL4+24* | 0.737 | 0.049 | 140 | -4.6194 | 2.281E-04 | *** |
| *JU1200 mL4 / JU751 mL4+24* | 0.996 | 0.093 | 140 | -0.0440 | 1.000E+00 | |
| *JU1200 mL4 / JU1200 YA* | 0.669 | 0.043 | 140 | -6.3033 | 9.882E-08 | *** |
| *JU1200 mL4 / JU751 YA* | 0.757 | 0.049 | 140 | -4.2803 | 8.825E-04 | *** |
| *JU1200 mL4 / JU1200 YA+24* | 0.928 | 0.079 | 140 | -0.8740 | 9.879E-01 | |
| *JU1200 mL4 / JU751 YA+24* | 1.204 | 0.113 | 140 | 1.9812 | 4.982E-01 | |
| *JU751 mL4 / JU1200 mL4+24* | 0.709 | 0.050 | 140 | -4.8499 | 8.691E-05 | *** |
| *JU751 mL4 / JU751 mL4+24* | 0.958 | 0.093 | 140 | -0.4403 | 9.998E-01 | |
| *JU751 mL4 / JU1200 YA* | 0.644 | 0.044 | 140 | -6.4087 | 5.812E-08 | *** |
| *JU751 mL4 / JU751 YA* | 0.729 | 0.051 | 140 | -4.5328 | 3.247E-04 | *** |
| *JU751 mL4 / JU1200 YA+24* | 0.893 | 0.080 | 140 | -1.2680 | 9.090E-01 | |
| *JU751 mL4 / JU751 YA+24* | 1.159 | 0.113 | 140 | 1.5161 | 7.976E-01 | |
| *JU1200 mL4+24 / JU751 mL4+24* | 1.351 | 0.106 | 140 | 3.8258 | 4.721E-03 | ** |
| *JU1200 mL4+24 / JU1200 YA* | 0.908 | 0.036 | 140 | -2.4195 | 2.403E-01 | |
| *JU1200 mL4+24 / JU751 YA* | 1.028 | 0.043 | 140 | 0.6544 | 9.980E-01 | |
| *JU1200 mL4+24 / JU1200 YA+24* | 1.259 | 0.087 | 140 | 3.3192 | 2.477E-02 | * |
| *JU1200 mL4+24 / JU751 YA+24* | 1.634 | 0.130 | 140 | 6.1786 | 1.842E-07 | *** |
| *JU751 mL4+24 / JU1200 YA* | 0.672 | 0.052 | 140 | -5.1787 | 2.072E-05 | *** |
| *JU751 mL4+24 / JU751 YA* | 0.760 | 0.059 | 140 | -3.5218 | 1.314E-02 | * |
| *JU751 mL4+24 / JU1200 YA+24* | 0.932 | 0.089 | 140 | -0.7387 | 9.956E-01 | |
| *JU751 mL4+24 / JU751 YA+24* | 1.209 | 0.125 | 140 | 1.8423 | 5.925E-01 | |
| *JU1200 YA / JU751 YA* | 1.131 | 0.043 | 140 | 3.2661 | 2.906E-02 | * |
| *JU1200 YA / JU1200 YA+24* | 1.386 | 0.093 | 140 | 4.8626 | 8.235E-05 | *** |
| *JU1200 YA / JU751 YA+24* | 1.799 | 0.139 | 140 | 7.5776 | 1.223E-10 | *** |
| *JU751 YA / JU1200 YA+24* | 1.225 | 0.084 | 140 | 2.9727 | 6.643E-02 | |
| *JU751 YA / JU751 YA+24* | 1.590 | 0.125 | 140 | 5.9056 | 7.024E-07 | *** |
| *JU1200 YA+24 / JU751 YA+24* | 1.298 | 0.125 | 140 | 2.7080 | 1.289E-01 | |

**Table S4 (accompanies Figure 2B). Statistical testing for differences in the number of dividing progenitor zone nuclei (EdU+, 15 min pulse) between JU1200 and JU751 at the mid L4 stage.** Data were fitted to a linear model with gaussian distribution: **EdU_Cells~ Strain** with a dispersion estimate (sigma^2) of 147 (A). The R software package, 'emmeans' was used to obtain estimated marginal means (B) and pairwise contrasts (C) with Tukey corrected p-values. For data representation, see Figure 2B.

**A) EdU+ PZ cells in JU1200 and JU751 at mL4: Gaussian conditional model**

| *Source* | *Estimate* | *SE* | *Z-value* | *Pr>\|z\|)* | |
|---|---|---|---|---|---|
| *Intercept* | 100.636 | 2.585 | 38.93 | <2e-16 | *** |
| *StrainJU751* | -15.956 | 3.545 | -4.50 | 6.75e-6 | *** |

**C) EdU+ PZ cells at mL4 in JU1200 and JU751 at mL4: Emmeans**

| *Strain* | *response* | *SE* | *df* |
|---|---|---|---|
| *JU1200* | 100.636 | 2.585 | 44 |
| *JU751* | 84.680 | 2.425 | 44 |

**D) EdU+ PZ cells at mL4 in JU1200 and JU751 at mL4: Contrasts**

| *Contrast* | *ratio* | *SE* | *df* | *t-ratio* | *p -value* | |
|---|---|---|---|---|---|---|
| *JU1200 / JU751* | 15.956 | 3.5447 | 44 | 4.502 | 4.90e-5 | *** |

**Table S5 (accompanies Figure 3A). Correction of block effect in the RIL data set.** RIL data were collected in six blocks (SB3-SB8), each of which contained both RIL parents. We performed a two-way ANOVA on the parental data alone to determine whether there was an effect of block on PZ Area. Formula: **Area~Line*Block.** Least squares means were used to derive a correction factor for each block by dividing the least squares mean of each block (SB4-SB8) by that of block SB3. Raw PZ area in pixels for each sample was multiplied by the correction factor to generate ‘Corr_PZ area’. The mean corrected PZ area (cMeanArea) for each RIL was used as the phenotype for QTL mapping (Data S1: Fig3A_mapping). Area in pixels was scaled to $\mu m^2/1000$ for presentation in Figures 3A, E, and H.

**A) ANOVA from only parental data from each block.**

| | df | Sum Sq | Mean Sq | F-value | Pr(>F) | |
|---|---|---|---|---|---|---|
| Line | 1 | 3987203584 | 3987203584 | 156.81 | 5.92E-30 | *** |
| Block | 5 | 3376974720 | 675394944 | 26.563 | 9.65E-23 | *** |
| Line:Block | 5 | 140124205 | 28024841 | 1.1022 | 0.3589 | ns |
| Residuals | 348 | 8848410248 | 25426466.23 | NA | NA | |

**B) Calculation of least squares means and correction factors.**

| Block | lsmean | SE | df | lower.CL | upper.CL | Corr_factor |
|---|---|---|---|---|---|---|
| SB3 | 33784.6 | 650.979598 | 348 | 32504.2506 | 35064.9494 | 1 |
| SB4 | 34671.3167 | 650.979598 | 348 | 33390.9672 | 35951.6661 | 1.02624618 |
| SB5 | 36650.4667 | 650.979598 | 348 | 35370.1172 | 37930.8161 | 1.0848276 |
| SB6 | 37485.3833 | 650.979598 | 348 | 36205.0339 | 38765.7328 | 1.10954054 |
| SB7 | 35956.65 | 650.979598 | 348 | 34676.3006 | 37236.9994 | 1.06429113 |
| SB8 | 43251.5333 | 650.979598 | 348 | 41971.1839 | 44531.8828 | 1.28021446 |

**Table S6 (accompanies Figure 3J) Statistical data for the multi-QTL model.** The fitqtl() function in R/qtl was used to generate model statistics for PZArea ~ chrII@pos37.0 + chrV@pos0.0.

**A) Full model result:**

Method: Haley-Knott regression
Model: normal phenotype
Number of observations: 70

| | df | SS | MS | LOD | %var | P value (Chi$^2$) | P value (F) | |
|---|---|---|---|---|---|---|---|---|
| Model | 2 | 2.0e8 | 1.0e8 | 5.85 | 31.96 | 1.4e-06 | 2.5e-06 | *** |
| Error | 67 | 4.3e8 | 6.4e6 | | | | | |
| Total | 69 | 6.3e7 | | | | | | |

**B) Drop one QTL at a time ANOVA table:**

| | df | Type III SS | LOD | %var | F value | P value (Chi$^2$) | P value (F) | |
|---|---|---|---|---|---|---|---|---|
| II@37.0 | 1 | 1.3e8 | 4.036 | 20.69 | 20.38 | 0 | 2.64e-05 | *** |
| V@0.0 | 1 | 8.6e7 | 2.794 | 13.73 | 13.52 | 0 | 0.000472 | *** |

**C) Estimated effects:**

| | estimate | SE | t |
|---|---|---|---|
| Intercept | 34477.6 | 303.9 | 113.437 |
| II@37.0 | -1371.9 | 303.9 | -4.514 |
| V@0.0 | -1119.6 | 304.5 | -3.677 |

**Table S7 (accompanies Figure 4C).** Statistical testing for differences in young adult progenitor zone size among the parental lines and JU1200 NILs. A) Data from nine experimental blocks were analyzed together. Raw PZ Area in pixels was fitted to a generalized linear mixed model with a negative binomial distribution and a log link, PZArea ~ Line*(1|Exp) with dispersion formula (~Line). B, C) The R software package, ‘emmeans’ was used to obtain estimated marginal means on the response scale and pairwise contrasts with Tukey corrected p-values. Area in pixels was scaled to $\mu m^2/1000$ for presentation in Figure 4C.

**A) PZ size across 9 experiments: Negative binomial conditional models**

Random effects:

| Groups | Source | Variance | Std.Dev |
|---|---|---|---|
| Exp | Intercept | 0.003648 | 0.0604 |

Number of obs: 1557, groups: Exp, 9

Main effects:

| Source | Estimate | SE | Z-value | Pr>\|z\|) | |
|---|---|---|---|---|---|
| Intercept | 11.45866 | 0.02123 | 539.7 | < 2e-16 | *** |
| LineJU751 | -0.22330 | 0.00979 | -22.8 | < 2e-16 | *** |
| LineNIC1697 | -0.05175 | 0.01054 | -4.9 | 9.04e-07 | *** |
| LineNIC1701 | -0.04231 | 0.01057 | -4.0 | 6.23e-05 | *** |

Dispersion model:

| Source | Estimate | SE | Z-value | Pr>\|z\|) | |
|---|---|---|---|---|---|
| Intercept | 3.87949 | 0.06531 | 59.40 | < 2e-16 | *** |
| LineJU751 | -0.19045 | 0.09405 | -2.02 | 0.0429 | * |
| LineNIC1697 | 0.16493 | 0.10868 | 1.52 | 0.1291 | |
| LineNIC1701 | 0.12415 | 0.10449 | 1.19 | 0.2348 | |

**B) PZ size in JU1200 NILs across 9 experiments: Emmeans**

| Strain | response | SE | df |
|---|---|---|---|
| JU1200 | 94717.8457 | 2010.82067 | 1548 |
| JU751 | 75762.3242 | 1625.40589 | 1548 |
| NIC1697 | 89940.653 | 1953.82761 | 1548 |
| NIC1701 | 90793.8034 | 1977.71553 | 1548 |
| JU1200 | 94717.8457 | 2010.82067 | 1548 |
| JU751 | 75762.3242 | 1625.40589 | 1548 |
| NIC1697 | 89940.653 | 1953.82761 | 1548 |
| NIC1701 | 90793.8034 | 1977.71553 | 1548 |
| JU1200 | 94717.8457 | 2010.82067 | 1548 |

**C) PZ size in JU1200 NILs across 9 experiments: Contrasts**

| Contrast | Ratio | SE | df | t-ratio | p -value | |
|---|---|---|---|---|---|---|
| JU1200 / JU751 | 1.2501972 | 0.01223992 | 1548 | 22.8082032 | 0 | *** |
| JU1200 / NIC1697 | 1.05311494 | 0.01109685 | 1548 | 4.91141477 | 5.96E-06 | *** |
| JU1200 / NIC1701 | 1.04321927 | 0.01102459 | 1548 | 4.00378144 | 0.00037965 | *** |
| JU751 / NIC1697 | 0.84235906 | 0.00922837 | 1548 | -15.65887 | 0 | *** |
| JU751 / NIC1701 | 0.83444378 | 0.00922435 | 1548 | -16.372524 | 0 | *** |
| NIC1697 / NIC1701 | 0.99060343 | 0.01155421 | 1548 | -0.8094267 | 0.85003654 | |

**Table S8 (accompanies Figure 4D). Statistical testing for differences in young adult Progenitor Zone size among the parental lines and JU751 NILs**. A) Raw data (PZ area in pixels) from one representative experiment of nine were fitted to a generalized linear model with a negative binomial distribution and a log link**, PZSize ~ Line**, with dispersion parameter of 60.2. B,C) The R software package, 'emmeans' was used to obtain estimated marginal means on the response scale and pairwise contrasts with Tukey corrected p-values. Area in pixels was scaled to $\mu m^2/1000$ for presentation in Figure 4D.

**A) PZ size in parental lines and JU751 NILs: Gaussian conditional model**

| Source | Estimate | SE | Z-value | Pr>\|z\|) | |
|---|---|---|---|---|---|
| Intercept | 10.81931 | 0.01922 | 562.8 | < 2e-16 | *** |
| LineJU751 | -0.21814 | 0.02719 | -8 | 1.03E-15 | *** |
| LineNIC1671 | -0.11557 | 0.02719 | -4.3 | 2.13E-05 | *** |
| LineNIC1672 | -0.07027 | 0.02719 | -2.6 | 0.00974 | ** |
| LineNIC1673 | -0.12449 | 0.02719 | -4.6 | 4.67E-06 | *** |
| LineNIC1675 | -0.0306 | 0.02719 | -1.1 | 0.26036 | |
| LineNIC1676 | -0.2371 | 0.02719 | -8.7 | < 2e-16 | *** |

**B) PZ size in parental lines and JU751 NILs: Emmeans**

| Line | response | SE | df |
|---|---|---|---|
| JU1200 | 49976.6843 | 960.742959 | 307 |
| JU751 | 40181.811 | 772.561924 | 307 |
| NIC1671 | 44522.2362 | 855.950461 | 307 |
| NIC1672 | 46585.1892 | 895.583778 | 307 |
| NIC1673 | 44126.8179 | 848.353619 | 307 |
| NIC1675 | 48470.5764 | 931.807047 | 307 |
| NIC1676 | 39427.0885 | 758.061678 | 307 |

**C) PZ size in parental lines and JU751 NILs: Contrasts**

| Contrast | ratio | SE | df | t-ratio | p -value | |
|---|---|---|---|---|---|---|
| JU1200 / JU751 | 1.24376386 | 0.03381619 | 307 | 8.02329592 | 1.63E-12 | *** |
| JU1200 / NIC1671 | 1.12251065 | 0.03051836 | 307 | 4.2507566 | 0.00055904 | *** |
| JU1200 / NIC1672 | 1.072802 | 0.02916645 | 307 | 2.58481887 | 0.13424479 | |
| JU1200 / NIC1673 | 1.13256942 | 0.03079193 | 307 | 4.57887184 | 0.00013742 | *** |
| JU1200 / NIC1675 | 1.03107262 | 0.02803161 | 307 | 1.12553123 | 0.91991339 | |
| JU1200 / NIC1676 | 1.26757228 | 0.03446375 | 307 | 8.72063522 | 1.17E-12 | *** |

| | | | | | | |
|---|---|---|---|---|---|---|
| *JU751 / NIC1671* | 0.90251107 | 0.02453891 | 307 | -3.7725581 | 0.00362754 | ** |
| *JU751 / NIC1672* | 0.86254476 | 0.02345188 | 307 | -5.4384959 | 2.28E-06 | *** |
| *JU751 / NIC1673* | 0.91059843 | 0.02475888 | 307 | -3.4444426 | 0.01150048 | * |
| *JU751 / NIC1675* | 0.82899388 | 0.02253939 | 307 | -6.8977734 | 6.35E-10 | *** |
| *JU751 / NIC1676* | 1.01914223 | 0.02771128 | 307 | 0.69734372 | 0.99268438 | |
| *NIC1671 / NIC1672* | 0.95571655 | 0.02598419 | 307 | -1.6659417 | 0.63930687 | |
| *NIC1671 / NIC1673* | 1.00896095 | 0.02743231 | 307 | 0.32811614 | 0.9998988 | |
| *NIC1671 / NIC1675* | 0.91854151 | 0.02497316 | 307 | -3.1252274 | 0.03167016 | * |
| *NIC1671 / NIC1676* | 1.12922962 | 0.03070351 | 307 | 4.46990136 | 0.00022128 | *** |
| *NIC1672 / NIC1673* | 1.0557115 | 0.02870295 | 307 | 1.99405778 | 0.42054372 | |
| *NIC1672 / NIC1675* | 0.96110244 | 0.0261299 | 307 | -1.4592878 | 0.76863618 | |
| *NIC1672 / NIC1676* | 1.18155286 | 0.03212567 | 307 | 6.13583829 | 5.46E-08 | *** |
| *NIC1673 / NIC1675* | 0.9103836 | 0.02475144 | 307 | -3.453343 | 0.01116249 | * |
| *NIC1673 / NIC1676* | 1.11920052 | 0.03043091 | 307 | 4.14178602 | 0.0008723 | *** |
| *NIC1675 / NIC1676* | 1.22937245 | 0.03342545 | 307 | 7.59511412 | 9.00E-12 | *** |

**Table S9 (accompanies Figure 5B). Statistical testing for differences in young adult Progenitor Zone size among the parents and ARLs**. A) Raw data (PZ area in pixels) were fitted to a generalized linear model with a negative binomial distribution and log link**, PZSize ~ Line**, with a dispersion parameter of 51.4. B,C) The R software package, 'emmeans' was used to obtain estimated marginal means on the response scale and pairwise contrasts with Tukey corrected p-values. Area in pixels was scaled to $\mu m^2/1000$ for presentation in Figure 5B.

**A) PZ size in parental lines and ARLs: Negative binomial conditional model**

| *Source* | *Estimate* | *SE* | *Z-value* | *Pr>\|z\|)* | |
|---|---|---|---|---|---|
| *Intercept* | 11.51157 | 0.01179 | 976.3 | <2e-16 | *** |
| *LineJU751* | -0.18316 | 0.01680 | -10.9 | <2e-16 | *** |
| *LineNIC1720* | -0.18556 | 0.01680 | -11.0 | <2e-16 | *** |
| *LineNIC1725* | -0.27452 | 0.01674 | -16.4 | <2e-16 | *** |

**B) PZ size in parental lines and ARLs: Emmeans**

| *Line* | *response* | *SE* | *df* |
|---|---|---|---|
| *JU1200* | 99854.4059 | 1177.32195 | 545 |
| *JU751* | 83142.2612 | 994.641831 | 545 |
| *NIC1720* | 82943.0346 | 992.260739 | 545 |
| *NIC1725* | 75883.0906 | 901.224303 | 545 |

**C) PZ size in parental lines and ARLs: Contrasts**

| *Contrast* | *ratio* | *SE* | *df* | *t-ratio* | *p -value* | |
|---|---|---|---|---|---|---|
| *JU1200 / JU751* | 1.20100662 | 0.02017297 | 545 | 10.9045148 | 3.89E-10 | *** |
| *JU1200 / NIC1720* | 1.2038914 | 0.02022145 | 545 | 11.0473327 | 3.89E-10 | *** |
| *JU1200 / NIC1725* | 1.31589798 | 0.0220217 | 545 | 16.4037903 | 3.89E-10 | *** |
| *JU751 / NIC1720* | 1.00240197 | 0.01695908 | 545 | 0.14180313 | 0.99898218 | |
| *JU751 / NIC1725* | 1.09566256 | 0.01846987 | 545 | 5.41957756 | 5.37E-07 | *** |
| *NIC1720 / NIC1725* | 1.09303712 | 0.01842564 | 545 | 5.27725369 | 1.13E-06 | *** |

**Table S10 (accompanies Figure 5D-F). Statistical testing for differences in DTC *lag-2* expression (smFISH) among the parents and ARLs**. A) Data were fitted to a generalized linear model with a negative binomial distribution and log link**, Puncta ~ Line*Stage**, with a dispersion parameter of 24.7. B, C) The R software package, 'emmeans' was used to obtain estimated marginal means on the response scale and pairwise contrasts with Tukey corrected p-values. For data representation, see Fig 5D-F.

**A) *lag-2* expression in parental lines and ARLs: Negative binomial conditional model**

| *Source* | *Estimate* | *SE* | *Z-value* | *Pr>\|z\|)* | |
|---|---|---|---|---|---|
| *Intercept* | 4.130 | 0.053 | 77.59 | <2e-16 | *** |
| *LineJU751* | -0.2347 | 0.077 | -3.06 | 0.0022 | ** |
| *LineNIC1720* | -0.0721 | 0.075 | 0.96 | 0.3354 | |
| *LineNIC1725* | -0.4986 | 0.079 | -6.34 | 2.31E-10 | *** |
| *StageeL3* | 0.6343 | 0.073 | 8.72 | <2e-16 | *** |
| *StagemL4* | 0.0563 | 0.075 | 0.75 | 0.4526 | |
| *LineJU751:StageeL3* | -0.0140 | 0.104 | -0.13 | 0.8937 | |
| *LineNIC1720:StageeL3* | -0.6532 | 0.104 | -6.27 | 3.6E-10 | *** |
| *LineNIC1725:StageeL3* | -0.0016 | 0.107 | 0.01 | 0.9882 | |
| *LineJU751:StagemL4* | 0.3358 | 0.107 | 3.15 | 0.0016 | ** |
| *LineNIC1720:StagemL4* | -0.0906 | 0.106 | -0.86 | 0.3921 | |
| *LineNIC1725:StagemL4* | 0.4617 | 0.109 | 4.23 | 2.39E-05 | *** |

**B) *lag-2* expression in parental lines and ARLs: Emmeans**

| Line | Stage | response | SE | df |
|---|---|---|---|---|
| *JU1200* | *mL2* | 62.150 | 3.308 | 226 |
| *JU751* | *mL2* | 49.150 | 2.712 | 226 |
| *NIC1720* | *mL2* | 66.800 | 3.520 | 226 |
| *NIC1725* | *mL2* | 37.750 | 2.186 | 226 |
| *JU1200* | *eL3* | 117.20 | 5.807 | 226 |
| *JU751* | *eL3* | 91.400 | 4.638 | 226 |
| *NIC1720* | *eL3* | 65.550 | 3.463 | 226 |
| *NIC1725* | *eL3* | 71.300 | 3.725 | 226 |
| *JU1200* | *mL4* | 65.750 | 3.472 | 226 |
| *JU751* | *mL4* | 72.750 | 3.791 | 226 |
| *NIC1720* | *mL4* | 64.550 | 3.417 | 226 |
| *NIC1725* | *mL4* | 63.368 | 3.451 | 226 |

**C) *lag-2* expression in parental lines and ARLs: Contrasts**

| Contrast | ratio | SE | df | t-ratio | p -value | |
|---|---|---|---|---|---|---|
| *JU1200 mL2 / JU751 mL2* | 1.264 | 0.097 | 226 | 3.061 | 0.0985 | |
| *JU1200 mL2 / NIC1720 mL2* | 0.9303 | 0.070 | 226 | -0.9633 | 0.9983 | |
| *JU1200 mL2 / NIC1725 mL2* | 1.6464 | 0.129 | 226 | 6.339 | 8.13E-08 | *** |
| *JU1200 mL2 / JU1200 eL3* | 0.5303 | 0.039 | 226 | -8.723 | 1.50E-13 | *** |
| *JU1200 mL2 / JU751 eL3* | 0.6800 | 0.050 | 226 | -5.245 | 2.30E-05 | *** |
| *JU1200 mL2 / NIC1720 eL3* | 0.9481 | 0.071 | 226 | -0.7102 | 0.9999 | |
| *JU1200 mL2 / NIC1725 eL3* | 0.8717 | 0.065 | 226 | -1.842 | 0.7932 | |
| *JU1200 mL2 / JU1200 mL4* | 0.9452 | 0.071 | 226 | -0.7510 | 0.9998 | |
| *JU1200 mL2 / JU751 mL4* | 0.8543 | 0.064 | 226 | -2.114 | 0.6132 | |
| *JU1200 mL2 / NIC1720 mL4* | 0.9628 | 0.072 | 226 | -0.5047 | 1 | |
| *JU1200 mL2 / NIC1725 mL4* | 0.9808 | 0.075 | 226 | -0.2550 | 1 | |
| *JU751 mL2 / NIC1720 mL2* | 0.7358 | 0.056 | 226 | -4.021 | 0.0044 | ** |
| *JU751 mL2 / NIC1725 mL2* | 1.302 | 0.104 | 226 | 3.299 | 0.0505 | |
| *JU751 mL2 / JU1200 eL3* | 0.4194 | 0.031 | 226 | -11.717 | 1.41E-14 | *** |
| *JU751 mL2 / JU751 eL3* | 0.5377 | 0.040 | 226 | -8.275 | 8.58E-13 | *** |
| *JU751 mL2 / NIC1720 eL3* | 0.7498 | 0.057 | 226 | -3.769 | 0.0111 | * |
| *JU751 mL2 / NIC1725 eL3* | 0.6893 | 0.052 | 226 | -4.895 | 0.0001 | *** |
| *JU751 mL2 / JU1200 mL4* | 0.7475 | 0.057 | 226 | -3.810 | 0.0096 | ** |
| *JU751 mL2 / JU751 mL4* | 0.6756 | 0.051 | 226 | -5.167 | 3.33E-05 | *** |
| *JU751 mL2 / NIC1720 mL4* | 0.7614 | 0.058 | 226 | -3.564 | 0.0221 | * |
| *JU751 mL2 / NIC1725 mL4* | 0.7756 | 0.060 | 226 | -3.277 | 0.0539 | |
| *NIC1720 mL2 / NIC1725 mL2* | 1.770 | 0.139 | 226 | 7.290 | 3.41E-10 | *** |
| *NIC1720 mL2 / JU1200 eL3* | 0.5700 | 0.041 | 226 | -7.772 | 1.78E-11 | *** |
| *NIC1720 mL2 / JU751 eL3* | 0.7308 | 0.053 | 226 | -4.286 | 0.0016 | ** |
| *NIC1720 mL2 / NIC1720 eL3* | 1.019 | 0.076 | 226 | 0.2532 | 1 | |
| *NIC1720 mL2 / NIC1725 eL3* | 0.9370 | 0.070 | 226 | -0.8786 | 0.9993 | |
| *NIC1720 mL2 / JU1200 mL4* | 1.016 | 0.076 | 226 | 0.2123 | 1 | |
| *NIC1720 mL2 / JU751 mL4* | 0.9182 | 0.068 | 226 | -1.151 | 0.9918 | |
| *NIC1720 mL2 / NIC1720 mL4* | 1.035 | 0.077 | 226 | 0.4587 | 1 | |
| *NIC1720 mL2 / NIC1725 mL4* | 1.054 | 0.080 | 226 | 0.6959 | 1 | |
| *NIC1725 mL2 / JU1200 eL3* | 0.3221 | 0.025 | 226 | -14.866 | 0 | *** |
| *NIC1725 mL2 / JU751 eL3* | 0.4130 | 0.032 | 226 | -11.485 | 2.78E-14 | *** |
| *NIC1725 mL2 / NIC1720 eL3* | 0.5759 | 0.045 | 226 | -7.040 | 1.50E-09 | *** |
| *NIC1725 mL2 / NIC1725 eL3* | 0.5295 | 0.041 | 226 | -8.154 | 1.72E-12 | *** |
| *NIC1725 mL2 / JU1200 mL4* | 0.5741 | 0.045 | 226 | -7.081 | 1.18E-09 | *** |
| *NIC1725 mL2 / JU751 mL4* | 0.5189 | 0.040 | 226 | -8.422 | 3.97E-13 | *** |
| *NIC1725 mL2 / NIC1720 mL4* | 0.5848 | 0.046 | 226 | -6.837 | 4.89E-09 | *** |
| *NIC1725 mL2 / NIC1725 mL4* | 0.5957 | 0.047 | 226 | -6.517 | 3.04E-08 | *** |
| *JU1200 eL3 / JU751 eL3* | 1.282 | 0.091 | 226 | 3.506 | 0.0267 | * |
| *JU1200 eL3 / NIC1720 eL3* | 1.788 | 0.129 | 226 | 8.023 | 3.80E-12 | *** |
| *JU1200 eL3 / NIC1725 eL3* | 1.644 | 0.118 | 226 | 6.902 | 3.36E-09 | *** |
| *JU1200 eL3 / JU1200 mL4* | 1.783 | 0.129 | 226 | 7.982 | 4.87E-12 | *** |
| *JU1200 eL3 / JU751 mL4* | 1.611 | 0.116 | 226 | 6.632 | 1.59E-08 | *** |
| *JU1200 eL3 / NIC1720 mL4* | 1.816 | 0.132 | 226 | 8.226 | 1.12E-12 | *** |
| *JU1200 eL3 / NIC1725 mL4* | 1.850 | 0.136 | 226 | 8.352 | 5.65E-13 | *** |
| *JU751 eL3 / NIC1720 eL3* | 1.394 | 0.102 | 226 | 4.538 | 0.0006 | *** |
| *JU751 eL3 / NIC1725 eL3* | 1.282 | 0.093 | 226 | 3.410 | 0.0361 | * |
| *JU751 eL3 / JU1200 mL4* | 1.390 | 0.102 | 226 | 4.497 | 0.0007 | *** |
| *JU751 eL3 / JU751 mL4* | 1.256 | 0.091 | 226 | 3.138 | 0.0801 | |
| *JU751 eL3 / NIC1720 mL4* | 1.416 | 0.104 | 226 | 4.743 | 0.0002 | *** |

| | | | | | | |
|---|---|---|---|---|---|---|
| *JU751 eL3 / NIC1725 mL4* | 1.442 | 0.107 | 226 | 4.921 | 0.0001 | *** |
| *NIC1720 eL3 / NIC1725 eL3* | 0.9194 | 0.068 | 226 | -1.132 | 0.9929 | |
| *NIC1720 eL3 / JU1200 mL4* | 0.9970 | 0.074 | 226 | -0.041 | 1 | |
| *NIC1720 eL3 / JU751 mL4* | 0.9010 | 0.067 | 226 | -1.404 | 0.9617 | |
| *NIC1720 eL3 / NIC1720 mL4* | 1.015 | 0.076 | 226 | 0.206 | 1 | |
| *NIC1720 eL3 / NIC1725 mL4* | 1.034 | 0.078 | 226 | 0.446 | 1 | |
| *NIC1725 eL3 / JU1200 mL4* | 1.084 | 0.081 | 226 | 1.091 | 0.9948 | |
| *NIC1725 eL3 / JU751 mL4* | 0.9801 | 0.072 | 226 | -0.273 | 1 | |
| *NIC1725 eL3 / NIC1720 mL4* | 1.105 | 0.082 | 226 | 1.337 | 0.9733 | |
| *NIC1725 eL3 / NIC1725 mL4* | 1.125 | 0.085 | 226 | 1.563 | 0.9205 | |
| *JU1200 mL4 / JU751 mL4* | 0.9038 | 0.067 | 226 | -1.364 | 0.9691 | |
| *JU1200 mL4 / NIC1720 mL4* | 1.019 | 0.076 | 226 | 0.2463 | 1 | |
| *JU1200 mL4 / NIC1725 mL4* | 1.038 | 0.079 | 226 | 0.4864 | 1 | |
| *JU751 mL4 / NIC1720 mL4* | 1.127 | 0.084 | 226 | 1.610 | 0.9039 | |
| *JU751 mL4 / NIC1725 mL4* | 1.148 | 0.087 | 226 | 1.832 | 0.7988 | |
| *NIC1720 mL4 / NIC1725 mL4* | 1.019 | 0.077 | 226 | 0.2433 | 1 | |

**Table S11 accompanies Figure 6. Statistical testing for differences in PZ area among the parents, NILs, and ARLs (“Interaction Data Set”) due to the genotypes of the background (BGND) the central portion of the chromosome II QTL (C2) and the chromosome V *lag-2* variant (C5*).*** A) Raw data (PZ area in pixels) were fitted to a generalized linear model with a negative binomial distribution and log link, **PZArea ~ BGND*C2 * C5 + BLOCK + BLOCK:BGND + BLOCK:C2** with a dispersion parameter of 44.55. Experimental block was included as fixed effect term with significant interactions in the model. Model selection was conducted using an ‘up-down’ approach to optimize both the Baysian Information Criterion (BIC) and the adjusted deviance accounted for in the model ($D^2$) using the R software package, modEvA (Márcia Barbosa et al., 2013). The package, ‘emmeans’ was used to obtain estimated marginal means on the response scale (B) and pairwise contrasts for the fixed effects (C) and interactions (D) with Tukey corrected p-values. Analysis was conducted on raw data (PZ area in pixels) and y-axes were scaled to ‘PZ Size’ in $\mu m^2/1000$ for data representation in Figure 6.

**Deviance Residuals:**

| Min | 1Q | Med | 3Q | Max |
|---|---|---|---|---|
| -3.748 | -0.73 | -0.031 | 0.656 | 3.033 |

**Null deviance:** 4233.7 on 2785 degrees of freedom
**Residual deviance:** 2796.5 on 2763 degrees of freedom
**AIC:** 60439
**BIC**: 60581.48
**$D^2$** = 0.3339762
**$p\chi^2$**= 0.3237755

**A) PZ Area from Interaction Data Set: Negative binomial conditional model**

| *Source* | *estimate* | *SE* | *Z-value* | *Pr>\|z\|)* | |
|---|---|---|---|---|---|
| *Intercept* | 11.35674 | 0.01681 | 675.679 | < 2e-16 | *** |
| *BGND JU751* | -0.12907 | 0.01872 | -6.893 | 5.44E-12 | *** |
| *C2 JU751* | -0.0436 | 0.0192 | -2.27 | 0.023188 | * |
| *C5 JU751* | -0.12523 | 0.01437 | -8.712 | < 2e-16 | *** |
| *BLOCK A3* | 0.18049 | 0.02117 | 8.524 | < 2e-16 | *** |
| *BLOCK B1* | 0.07787 | 0.02053 | 3.792 | 0.000149 | *** |
| *BLOCK B2* | 0.13552 | 0.02021 | 6.706 | 1.99E-11 | *** |
| *BLOCK N3* | 0.19258 | 0.02371 | 8.122 | 4.57E-16 | *** |
| *BLOCK N5* | 0.07279 | 0.02559 | 2.845 | 0.004442 | ** |
| *BGND JU751:C2 JU751* | -0.06228 | 0.01855 | -3.357 | 0.000788 | *** |
| *BGND JU751:C5 JU751* | 0.15208 | 0.01932 | 7.872 | 3.48E-15 | *** |
| *C2 JU751:C5 JU751* | 0.06673 | 0.01906 | 3.501 | 0.000463 | *** |
| *BGND JU751:BLOCK A3* | -0.17217 | 0.02511 | -6.857 | 7.02E-12 | *** |
| *BGND JU751:BLOCK B1* | -0.02981 | 0.02249 | -1.326 | 0.184893 | |
| *BGND JU751:BLOCK B2* | -0.04634 | 0.02165 | -2.14 | 0.03235 | * |
| *BGND JU751:BLOCK N3* | -0.02818 | 0.02474 | -1.139 | 0.254816 | |
| *BGND JU751:BLOCK N5* | -0.13209 | 0.02683 | -4.923 | 8.52E-07 | *** |
| *C2 JU751:BLOCKA3* | -0.06427 | 0.0235 | -2.735 | 0.006239 | ** |
| *C2 JU751:BLOCKB1* | 0.00655 | 0.02093 | 0.313 | 0.754354 | |
| *C2 JU751:BLOCKB2* | 0.04348 | 0.02016 | 2.157 | 0.031009 | * |
| *C2 JU751:BLOCKN3* | -0.05689 | 0.02418 | -2.353 | 0.018624 | * |
| *C2 JU751:BLOCKN5* | -0.02327 | 0.02613 | -0.891 | 0.373151 | |
| *BGND JU751:C2 JU751:C5 JU751* | -0.01975 | 0.02669 | -0.74 | 0.459272 | |

**B) PZ Area from Interaction Data Set: Emmeans (BGND, C2, C5)**

| BGND | C2 | C5 | response | SE | df |
|---|---|---|---|---|---|
| JU1200 | JU1200 | JU1200 | 95474.4935 | 786.532429 | Inf |
| JU751 | JU1200 | JU1200 | 78390.0105 | 701.590905 | Inf |
| JU1200 | JU751 | JU1200 | 89974.616 | 819.559659 | Inf |
| JU751 | JU751 | JU1200 | 69413.5523 | 773.624993 | Inf |
| JU1200 | JU1200 | JU751 | 84236.665 | 1058.236 | Inf |
| JU751 | JU1200 | JU751 | 80523.564 | 647.430365 | Inf |
| JU1200 | JU751 | JU751 | 84862.5242 | 779.008679 | Inf |
| JU751 | JU751 | JU751 | 74732.61 | 589.913664 | Inf |

**C) PZ Area from Interaction Data Set: Emmeans (BGND, C2)**

| BGND | C2 | response | SE | df |
|---|---|---|---|---|
| JU1200 | JU1200 | 89679.7241 | 701.530732 | Inf |
| JU751 | JU1200 | 79449.6258 | 433.220653 | Inf |
| JU1200 | JU751 | 87381.1938 | 508.136821 | Inf |
| JU751 | JU751 | 72023.9955 | 497.934192 | Inf |

**D) PZ Area from Interaction Data Set: Emmeans (BGND, C5)**

| BGND | C5 | response | SE | df |
|---|---|---|---|---|
| JU1200 | JU1200 | 92683.7682 | 562.992739 | Inf |
| JU751 | JU1200 | 73765.3652 | 551.563165 | Inf |
| JU1200 | JU751 | 84549.0155 | 700.082394 | Inf |
| JU751 | JU751 | 77574.0685 | 439.703409 | Inf |

**E) PZ Area from Interaction Data Set: Emmeans (C2, C5)**

| C2 | C5 | response | SE | df |
|---|---|---|---|---|
| JU1200 | JU1200 | 86511.54 | 527.213468 | Inf |
| JU751 | JU1200 | 79028.2083 | 554.518207 | Inf |
| JU1200 | JU751 | 82359.1919 | 612.814813 | Inf |
| JU751 | JU751 | 79636.6619 | 479.872666 | Inf |

**F) PZ Area from Interaction Data Set: Contrasts**

| Contrast (BGND C2 C5 / BGND C2 C5) | ratio | SE | df | t-ratio | p-value | |
|---|---|---|---|---|---|---|
| JU1200 JU1200 JU1200 / JU751 JU1200 JU1200 | 1.2179 | 0.0148 | Inf | 16.2405 | 0 | *** |
| JU1200 JU1200 JU1200 / JU1200 JU751 JU1200 | 1.0611 | 0.0132 | Inf | 4.7798 | 4.80E-05 | *** |
| JU1200 JU1200 JU1200 / JU751 JU751 JU1200 | 1.3754 | 0.0195 | Inf | 22.5214 | 0 | *** |
| JU1200 JU1200 JU1200 / JU1200 JU1200 JU751 | 1.1334 | 0.0163 | Inf | 8.7124 | 6.54E-14 | *** |
| JU1200 JU1200 JU1200 / JU751 JU1200 JU751 | 1.1857 | 0.0137 | Inf | 14.7176 | 0 | *** |
| JU1200 JU1200 JU1200 / JU1200 JU751 JU751 | 1.1250 | 0.0137 | Inf | 9.6894 | 9.66E-14 | *** |
| JU1200 JU1200 JU1200 / JU751 JU751 JU751 | 1.2775 | 0.0147 | Inf | 21.3024 | 0 | *** |
| JU751 JU1200 JU1200 / JU1200 JU751 JU1200 | 0.8712 | 0.0110 | Inf | -10.919 | 7.77E-14 | *** |
| JU751 JU1200 JU1200 / JU751 JU751 JU1200 | 1.1293 | 0.0154 | Inf | 8.9414 | 5.58E-14 | *** |
| JU751 JU1200 JU1200 / JU1200 JU1200 JU751 | 0.9306 | 0.0144 | Inf | -4.6344 | 9.72E-05 | *** |
| JU751 JU1200 JU1200 / JU751 JU1200 JU751 | 0.9735 | 0.0127 | Inf | -2.0561 | 0.4438 | |
| JU751 JU1200 JU1200 / JU1200 JU751 JU751 | 0.9237 | 0.0119 | Inf | -6.1626 | 2.00E-08 | *** |
| JU751 JU1200 JU1200 / JU751 JU751 JU751 | 1.0489 | 0.0126 | Inf | 3.9874 | 0.0017 | ** |
| JU1200 JU751 JU1200 / JU751 JU751 JU1200 | 1.2962 | 0.0191 | Inf | 17.5948 | 0 | *** |
| JU1200 JU751 JU1200 / JU1200 JU1200 JU751 | 1.0681 | 0.0172 | Inf | 4.0975 | 0.0012 | ** |
| JU1200 JU751 JU1200 / JU751 JU1200 JU751 | 1.1174 | 0.0137 | Inf | 9.0820 | 1.04E-13 | *** |
| JU1200 JU751 JU1200 / JU1200 JU751 JU751 | 1.0602 | 0.0150 | Inf | 4.1444 | 0.0009 | *** |
| JU1200 JU751 JU1200 / JU751 JU751 JU751 | 1.2040 | 0.0145 | Inf | 15.4204 | 0 | *** |
| JU751 JU751 JU1200 / JU1200 JU1200 JU751 | 0.8240 | 0.0143 | Inf | -11.159 | 2.49E-14 | *** |
| JU751 JU751 JU1200 / JU751 JU1200 JU751 | 0.8620 | 0.0121 | Inf | -10.544 | 7.37E-14 | *** |
| JU751 JU751 JU1200 / JU1200 JU751 JU751 | 0.8180 | 0.0116 | Inf | -14.157 | 0 | *** |
| JU751 JU751 JU1200 / JU751 JU751 JU751 | 0.9288 | 0.0125 | Inf | -5.4750 | 1.22E-06 | *** |
| JU1200 JU1200 JU751 / JU751 JU1200 JU751 | 1.0461 | 0.0156 | Inf | 3.0156 | 0.0523 | . |
| JU1200 JU1200 JU751 / JU1200 JU751 JU751 | 0.9926 | 0.0144 | Inf | -0.5109 | 0.9996 | |
| JU1200 JU1200 JU751 / JU751 JU751 JU751 | 1.1272 | 0.0169 | Inf | 7.9853 | 1.48E-13 | *** |
| JU751 JU1200 JU751 / JU1200 JU751 JU751 | 0.9489 | 0.0116 | Inf | -4.2780 | 0.0005 | *** |
| JU751 JU1200 JU751 / JU751 JU751 JU751 | 1.0775 | 0.0121 | Inf | 6.6648 | 7.42E-10 | *** |
| JU1200 JU751 JU751 / JU751 JU751 JU751 | 1.1355 | 0.0138 | Inf | 10.4520 | 7.75E-14 | *** |

**G) PZ Area from Interaction Data Set: Contrasts of Interactions**

| model term | by | df1 | df2 | F-ratio | p -value | |
|---|---|---|---|---|---|---|
| BGND:C2 | null | 1 | Inf | 32.599 | <0.0001 | *** |
| BGND:C5 | null | 1 | Inf | 96.984 | <0.0001 | *** |
| C2:C5 | null | 1 | Inf | 15.452 | <0.0001 | *** |

| | | | | | | |
|---|---|---|---|---|---|---|
| *BGND:C2* | *C5=JU1200* | 1 | Inf | 11.269 | 0.0008 | *** |
| *BGND:C2* | *C5=JU751* | 1 | Inf | 20.304 | <0.0001 | *** |
| *BGND:C5* | C2=*JU1200* | 1 | Inf | 61.967 | <0.0001 | *** |
| *BGND:C5* | C2=*JU751* | 1 | Inf | 43.773 | <0.0001 | *** |
| *C2:C5* | *BGND=JU1200* | 1 | Inf | 12.258 | 0.0005 | *** |
| *C2:C5* | *BGND=JU751* | 1 | Inf | 6.877 | 0.0087 | ** |
| *BGND:C2:C5* | *null* | 1 | Inf | 0.548 | 0.4593 | |

**Table S12 accompanies Figure S4. Statistical testing for differences in DTC *lag-2* expression in JU1200 and JU751 at mid L3 and young adult**. A) Data were fitted to a generalized linear model with a negative binomial distribution and log link**, Puncta ~ Line*Stage** with a dispersion parameter of 66.4. B,C) The R software package, 'emmeans' was used to obtain estimated marginal means on the response scale and pairwise contrasts with Tukey corrected p-values. For data representation, see Figure S4.

**A) Number of *lag-2* puncta in parental lines: Negative binomial conditional model**

| *Source* | *Estimate* | *SE* | *Z-value* | *Pr>\|z\|)* | |
|---|---|---|---|---|---|
| *Intercept* | 4.77153 | 0.03429 | 139.15 | <2e-16 | *** |
| *LineJU751* | -0.13389 | 0.04912 | -2.73 | 0.00641 | ** |
| *StageYA* | -0.77793 | 0.05339 | -14.57 | <2e-16 | *** |
| *LineJU751:StageYA* | -0.16475 | 0.07672 | 2.18 | 0.02956 | * |

**B) Number of *lag-2* puncta in parental lines: Emmeans**

| *Line* | *Stage* | *response* | *SE* | *df* |
|---|---|---|---|---|
| *JU1200* | *mL3* | 118.10 | 4.050 | 75 |
| *JU751* | *mL3* | 103.30 | 3.633 | 75 |
| *JU1200* | *YA* | 54.25 | 2.220 | 75 |
| *JU751* | *YA* | 55.95 | 2.270 | 75 |

**C) Number of *lag-2* puncta in parental lines: Contrasts**

| *Contrast* | *ratio* | *SE* | *df* | *t-ratio* | *p -value* | |
|---|---|---|---|---|---|---|
| *JU1200 mL3 / JU751 mL3* | 1.143 | 0.056 | 75 | 2.726 | 0.0390 | * |
| *JU1200 mL3 / JU1200 YA* | 2.177 | 0.116 | 75 | 14.572 | 0 | *** |
| *JU1200 mL3 / JU751 YA* | 2.111 | 0.112 | 75 | 14.063 | 0 | *** |
| *JU751 mL3 / JU1200 YA* | 1.904 | 0.103 | 75 | 11.937 | 0 | *** |
| *JU751 mL3 / JU751 YA* | 1.846 | 0.099 | 75 | 11.420 | 0 | *** |
| *JU1200 YA / JU751 YA* | 0.970 | 0.056 | 75 | -0.535 | 0.9501 | |

**Table S13. Oligonucleotides used in this study.**

| Names | Sequence (5'-3') | Purpose | Notes |
|---|---|---|---|
| plag-2 repair F1 | CAAATAGAGGGGTGTCGAAATGATAAATGTTGCA AGTGAAGTGCAGATGG | to generate CRISPR repair template | amplifies a 278bp fragment in JU1200 used as a repair template to replace the deleted 148bp sequence upstream of *lag-2* in JU751 |
| plag-2 repair R1 | GTTCAATGCACCGTCAAACCATATGAAACTTTATG GGCCGCTG | | |
| JU751 chrV sgRNA4 | AGACAGUCAGCGGCCCAUAA+GUUUUAGAGCUA UGCUGUUUUG | to generate allelic replacement lines using CRISPR | cuts downstream of deletion sequence |
| JU751 chrV sgRNA1 | AAUGAUAAAUGUUGCAAGUG+GUUUUAGAGCUA UGCUGUUUUG | | cuts upstream of deletion sequence |
| plag-2_ssODN_repair template | CAAATAGAGGGGTGTCGAAATGATAAATGTTGCA AGTGAAGTGCAGATGGCAATGAAATTTATATATG TTTGGGGTGTCTAGACAGTCAGCGGCCCATAAA GTTTCATATGGTTTGACGGTGCATTGAACT | to generate allelic replacement lines using CRISPR | repair template creates the 148bp *lag-2* deletion in JU1200 |
| dpy-10_sgRNA | GCUACCAUAGGCACCACGAG+GUUUUAGAGCUA UGCUGUUUUG | co-CRISPR guide RNA | cuts in *dpy-10* |
| dpy-10_ssODN_repair | CACTTGAACTTCAATACGGCAAGATGAGAATGAC TGGAAACCGTACCGCATGCGGTGCCTATGGTAG CGGAGCTT CACATGGCTTCAGACCAACAGCCTAT | co-CRISPR repair template | repair template introduces missense mutation (*cn64*) in *dpy-10* at cut-site |
| JU751cV del3.2 fwd1 | TGACTCTATTGCTCCTAATCCC | to genotype *plag-2* deletion region, not all combinations generate bands due to polyG repeat. See .ape file for JU751 and JU1200 versions of *plag-2* deletion region. | |
| JU751cV del3.2 rev1 | GAACAAAGGCATGAAGAGGG | | |
| JU751cV del3.2 fwd2 | CAAGAATAACAACAGGTGACTCG | | |
| JU751cV del3.2 rev2 | GTATTTCCAACTCTTCCCGTCTC | | |
| JU751cV del3.2 F3 | GGGTGTCGAAATGATAAATGTTGC | | |
| JU751cV del3-2 R3 | ACAAGAATGGAATGAACCCTTATGG | | |
| JU751cV del3-2 F5 | CAAGCAGAGCGAGAATTTGAG | | |
| JU751cV del3-2 R5 | CTCAAATTCTCGCTCTGCTTG | | |
| JU751cV del3-2 R4 | GGCAACTTTTCACTGATAATTTCAG | | |
| JU751cII del84 fwd1 | TAAGGCAGGCATGGGAACAG | genotype chrII QTL | amplifies INDEL 3916232-3916279 |
| JU751cII del84 rev1 | GAGGAAGAACACGAGTCAGGAAA | | |
| JU751cII del1 fwd1 | GCAGCAACACATCCCAGAAAG | genotype chrII QTL | amplifies INDEL 4187224-4187277 |
| JU751cII del1 rev1 | CTCCATACCTGCCTGCCTAC | | |
| JU751cII del3 fwd1 | GGTGGAGGTGAATGGAGAATTGC | genotype chrII QTL | amplifies INDEL 4265899-4266032 |
| JU751cII del3 rev1 | GGTAGGTAGGCAGGTAGGCA | | |
| JU751cII del12 fwd1 | CCGGAGTTCGTTGTTATGGTTGTAG | genotype chrII QTL | amplifies INDEL 4783409-4783461 |
| JU751cII del12 rev1 | GAGTAGAGCTGATACGGTGGCT | | |
| JU751cII del20 fwd1 | GGGACCTCCTACTTTCTGAATCG | genotype chrII QTL | amplifies INDEL 5968491-5968491 |
| JU751cII del20 rev1 | CGTGCCTACAGCTCCTACTC | | |
| JU751cII del27 fwd1 | AACCTGGGCACGGAAACTTG | genotype chrII QTL | amplifies INDEL 8429315-8429315 |
| JU751cII del27 rev1 | CTTCACCTTCCTGAGCGCATT | | |
| JU751cII del30 fwd1 | ATGCCTACCTTCTCCCACTTG | genotype chrII QTL | amplifies INDEL 9269605-9269650 |
| JU751cII del30 rev1 | GTTGCTCGTTCTCGGTTTGGATA | | |
| JU751cII del39 fwd1 | CAACACAAACAGCCACAAGATCAG | genotype chrII QTL | amplifies INDEL 10433097-10433230 |
| JU751cII del39 rev1 | GTGAAGAGGAGGAGGTGGATG | | |

| | | | |
|---|---|---|---|
| JU751cII del46 fwd1 | GTTACCTGTGCGTCGTTTGTC | genotype chrII QTL | amplifies INDEL 11511577-11511708 |
| JU751cII del46 rev1 | GACCTCTATTCTCCCGCCATT | | |
| JU751cII del49 fwd1 | GAGTGAGAGAGTGGGCGAAC | genotype chrII QTL | amplifies INDEL 11736927-11737094 |
| JU751cII del49 rev1 | ACGACGAGAGAGAGAGGATTCTG | | |
| JU751cII del68 fwd1 | AAACCGCCAGCTCCACCAAG | genotype chrII QTL | amplifies INDEL 12829541-12829633 |
| JU751cII del68 rev1 | GAGACCAATGCCGGGACCAT | | |
| JU751cII del72 fwd1 | CGCATCAGAACGGAGCACAG | genotype chrII QTL | amplifies INDEL 13339252-13339564 |
| JU751cII del72 rev1 | GGTTTCCTGAGCACTGACTTGTTG | | |
| JU751cII del79 fwd1 | TTTCTCCTTGTGCTGGCCTG | genotype chrII QTL | amplifies INDEL 13751711-13751802 |
| JU751cII del79 rev1 | CATGTGGCCGTGGAAGAGAC | | |
| JU751cII del81 fwd1 | AGGAACTGGAACCAAGAGAAGAGG | genotype chrII QTL | amplifies INDEL 13915292-13915529 |
| JU751cII del81 rev1 | CAAGAAATAGGCGGCGGAGC | | |
| lag-2_1 | GTAAGAGGAAGTAAGCGATC | smFISH probe (Stellaris ®) | detects *lag-2* mRNA |
| lag-2_2 | TCTTGATGGACTTCTACTCG | smFISH probe (Stellaris ®) | detects *lag-2* mRNA |
| lag-2_3 | ACTCGCTTCGAGGAGATGAA | smFISH probe (Stellaris ®) | detects *lag-2* mRNA |
| lag-2_4 | TGTGACAATTTCGAAGCGGA | smFISH probe (Stellaris ®) | detects *lag-2* mRNA |
| lag-2_5 | AAATGTGACTGGACGATTCG | smFISH probe (Stellaris ®) | detects *lag-2* mRNA |
| lag-2_6 | AGGATGATGTTGGTTTTGGG | smFISH probe (Stellaris ®) | detects *lag-2* mRNA |
| lag-2_7 | AAGTTGAAGACCGGGTTGAA | smFISH probe (Stellaris ®) | detects *lag-2* mRNA |
| lag-2_8 | AACGGCTGCACAAGTTGAAT | smFISH probe (Stellaris ®) | detects *lag-2* mRNA |
| lag-2_9 | TTTCGATAGATCCGATCACC | smFISH probe (Stellaris ®) | detects *lag-2* mRNA |
| lag-2_10 | TTGGTGCCCGAAAACTGAAC | smFISH probe (Stellaris ®) | detects *lag-2* mRNA |
| lag-2_11 | TGAACGTGTCGTTGATCCAC | smFISH probe (Stellaris ®) | detects *lag-2* mRNA |
| lag-2_12 | ACGAAATTCCAGATGTCGTC | smFISH probe (Stellaris ®) | detects *lag-2* mRNA |
| lag-2_13 | ATAGTTGCGAGCACATGTGA | smFISH probe (Stellaris ®) | detects *lag-2* mRNA |
| lag-2_14 | AAGTTTTCGCATCGATTCCC | smFISH probe (Stellaris ®) | detects *lag-2* mRNA |
| lag-2_15 | TTCGCCAAGTGTGCATCACA | smFISH probe (Stellaris ®) | detects *lag-2* mRNA |
| lag-2_16 | ATGGCGTCGCATCTTTTCCT | smFISH probe (Stellaris ®) | detects *lag-2* mRNA |
| lag-2_17 | ACAATGTGGTCCCATCCATC | smFISH probe (Stellaris ®) | detects *lag-2* mRNA |
| lag-2_18 | TCGTTCTCGCATGAGCACTT | smFISH probe (Stellaris ®) | detects *lag-2* mRNA |
| lag-2_19 | TGGATGAATCATCGACGAGA | smFISH probe (Stellaris ®) | detects *lag-2* mRNA |
| lag-2_20 | CGATGTTTGATTTGGCTGAC | smFISH probe (Stellaris ®) | detects *lag-2* mRNA |
| lag-2_21 | GCAGATCAACTGTTCATTGC | smFISH probe (Stellaris ®) | detects *lag-2* mRNA |
| lag-2_22 | AAAATTTCGCAACGAGTGCC | smFISH probe (Stellaris ®) | detects *lag-2* mRNA |
| lag-2_23 | TCAGCTGGAACTGATTGAAG | smFISH probe (Stellaris ®) | detects *lag-2* mRNA |
| lag-2_24 | TTTGACACTGCAAGCATCTG | smFISH probe (Stellaris ®) | detects *lag-2* mRNA |
| lag-2_25 | AAGCATTTGGCGCCATTCAA | smFISH probe (Stellaris ®) | detects *lag-2* mRNA |
| lag-2_26 | AGAAGACTTTCGGACCGTTT | smFISH probe (Stellaris ®) | detects *lag-2* mRNA |
| lag-2_27 | ATTCACCGATGAAGCCGACA | smFISH probe (Stellaris ®) | detects *lag-2* mRNA |
| lag-2_28 | TGGTGAGTGAGATTTCACAG | smFISH probe (Stellaris ®) | detects *lag-2* mRNA |
| lag-2_29 | TAATCTCCACGGTAGTTGGG | smFISH probe (Stellaris ®) | detects *lag-2* mRNA |
| lag-2_30 | AGCAACTGTGATGTAGACGG | smFISH probe (Stellaris ®) | detects *lag-2* mRNA |
| lag-2_31 | GATGGAGAAGATCACGAAGA | smFISH probe (Stellaris ®) | detects *lag-2* mRNA |

| | | | |
|---|---|---|---|
| lag-2_32 | GTACTTGAAGCATCCGATGA | smFISH probe (Stellaris ®) | detects *lag-2* mRNA |
| lag-2_33 | CTGTCTCATCGGCTTGAACT | smFISH probe (Stellaris ®) | detects *lag-2* mRNA |
| lag-2_34 | ATCTTGTAGGGCTCTGGTAC | smFISH probe (Stellaris ®) | detects *lag-2* mRNA |
| lag-2_35 | AGCATCGACTTTGTCTCAGG | smFISH probe (Stellaris ®) | detects *lag-2* mRNA |
| lag-2_36 | TCTGAAGCTTCCGGATCGAT | smFISH probe (Stellaris ®) | detects *lag-2* mRNA |
| lag-2_37 | TGGTGAACACCTTCTTCTGA | smFISH probe (Stellaris ®) | detects *lag-2* mRNA |
| lag-2_38 | AATCTTCTGCACACTGCCCT | smFISH probe (Stellaris ®) | detects *lag-2* mRNA |
| lag-2_39 | GGTATACCTCACTTCCTCAT | smFISH probe (Stellaris ®) | detects *lag-2* mRNA |
| lag-2_40 | TTCATACTTCCTTGGAGCAC | smFISH probe (Stellaris ®) | detects *lag-2* mRNA |
| lag-2_41 | AACGGCATACTCATTATTCG | smFISH probe (Stellaris ®) | detects *lag-2* mRNA |
| lag-2_42 | CGGTGGAGTAGACTTTTGAA | smFISH probe (Stellaris ®) | detects *lag-2* mRNA |
| lag-2_43 | ACGGTGGACTTAAAGATGGT | smFISH probe (Stellaris ®) | detects *lag-2* mRNA |
| lag-2_44 | CATAGTGACAGGCTGGAATA | smFISH probe (Stellaris ®) | detects *lag-2* mRNA |

**Table S14. Strains used in this study**

| Strain | Species | Genotype | Description | SOURCE |
|---|---|---|---|---|
| AF16, | *C. briggsae* | | wild isolate | *Caenorhabditis* Genetics Center (https://cgc.umn.edu) |
| CB4856 | *C. elegans* | | wild isolate | *C. elegans* Natural Diversity Resource (https://elegansvariation.org) |
| CX11262 | *C. elegans* | | wild isolate | *C. elegans* Natural Diversity Resource (https://elegansvariation.org) |
| HK104 | *C. briggsae* | | wild isolate | *Caenorhabditis* Genetics Center (https://cgc.umn.edu) |
| JU576 | *C. elegans* | | wild isolate | M.-A. Félix[74] |
| JU577 | *C. elegans* | | wild isolate | M.-A. Félix[74] |
| JU584 | *C. elegans* | | wild isolate | M.-A. Félix[74] |
| JU636 | *C. elegans* | | wild isolate | M.-A. Félix[74] |
| JU637 | *C. elegans* | | wild isolate | M.-A. Félix[74] |
| JU644 | *C. elegans* | | wild isolate | M.-A. Félix[74] |
| JU656 | *C. elegans* | | wild isolate | M.-A. Félix[74] |
| JU660 | *C. elegans* | | wild isolate | M.-A. Félix[74] |
| JU751 | *C. elegans* | | wild isolate | M.-A. Félix[74] |
| JU752 | *C. elegans* | | wild isolate | M.-A. Félix[74] |
| JU753 | *C. elegans* | | wild isolate | M.-A. Félix[74] |
| JU754 | *C. elegans* | | wild isolate | M.-A. Félix[74] |
| JU755 | *C. elegans* | | wild isolate | M.-A. Félix[74] |
| JU756 | *C. elegans* | | wild isolate | M.-A. Félix[74] |
| JU775 | *C. elegans* | | wild isolate | *C. elegans* Natural Diversity Resource (https://elegansvariation.org) |
| JU815 | *C. elegans* | | wild isolate | M.-A. Félix[74] |
| JU816 | *C. elegans* | | wild isolate | M.-A. Félix[74] |
| JU817 | *C. elegans* | | wild isolate | M.-A. Félix[74] |
| JU818 | *C. elegans* | | wild isolate | M.-A. Félix[74] |
| JU821 | *C. elegans* | | wild isolate | M.-A. Félix[74] |
| JU1200 | *C. elegans* | | wild isolate | *C. elegans* Natural Diversity Resource (https://elegansvariation.org) |
| JU1491 | *C. elegans* | | wild isolate | *C. elegans* Natural Diversity Resource (https://elegansvariation.org) |
| N2 | *C. elegans* | | Wild isolate, lab reference strain | *Caenorhabditis* Genetics Center (https://cgc.umn.edu) |
| NIC601 | *C. elegans* | | F2 RIL(1) JU751 x JU1200 | C. Braendle[70] |
| NIC606 | *C. elegans* | | F2 RIL(6) JU751 x JU1200 | C. Braendle[70] |
| NIC610 | *C. elegans* | | F2 RIL(10) JU751 x JU1200 | C. Braendle[70] |
| NIC617 | *C. elegans* | | F2 RIL(17) JU751 x JU1200 | C. Braendle[70] |

| | | | | |
|---|---|---|---|---|
| NIC619 | *C. elegans* | | F2 RIL(19) JU751 x JU1200 | C. Braendle[70] |
| NIC621 | *C. elegans* | | F2 RIL(21) JU751 x JU1200 | C. Braendle[70] |
| NIC623 | *C. elegans* | | F2 RIL(23) JU751 x JU1200 | C. Braendle[70] |
| NIC624 | *C. elegans* | | F2 RIL(24) JU751 x JU1200 | C. Braendle[70] |
| NIC625 | *C. elegans* | | F2 RIL(25) JU751 x JU1200 | C. Braendle[70] |
| NIC626 | *C. elegans* | | F2 RIL(26) JU751 x JU1200 | C. Braendle[70] |
| NIC628 | *C. elegans* | | F2 RIL(28) JU751 x JU1200 | C. Braendle[70] |
| NIC630 | *C. elegans* | | F2 RIL(30) JU751 x JU1200 | C. Braendle[70] |
| NIC631 | *C. elegans* | | F2 RIL(31) JU751 x JU1200 | C. Braendle[70] |
| NIC633 | *C. elegans* | | F2 RIL(33) JU751 x JU1200 | C. Braendle[70] |
| NIC636 | *C. elegans* | | F2 RIL(36) JU751 x JU1200 | C. Braendle[70] |
| NIC640 | *C. elegans* | | F2 RIL(40) JU751 x JU1200 | C. Braendle[70] |
| NIC643 | *C. elegans* | | F2 RIL(43) JU751 x JU1200 | C. Braendle[70] |
| NIC646 | *C. elegans* | | F2 RIL(46) JU751 x JU1200 | C. Braendle[70] |
| NIC647 | *C. elegans* | | F2 RIL(47) JU751 x JU1200 | C. Braendle[70] |
| NIC649 | *C. elegans* | | F2 RIL(49) JU751 x JU1200 | C. Braendle[70] |
| NIC650 | *C. elegans* | | F2 RIL(50) JU751 x JU1200 | C. Braendle[70] |
| NIC653 | *C. elegans* | | F2 RIL(53) JU751 x JU1200 | C. Braendle[70] |
| NIC654 | *C. elegans* | | F2 RIL(54) JU751 x JU1200 | C. Braendle[70] |
| NIC655, | *C. elegans* | | F2 RIL(55) JU751 x JU1200 | C. Braendle[70] |
| NIC660 | *C. elegans* | | F2 RIL(60) JU751 x JU1200 | C. Braendle[70] |
| NIC661 | *C. elegans* | | F2 RIL(61) JU751 x JU1200 | C. Braendle[70] |
| NIC662 | *C. elegans* | | F2 RIL(62) JU751 x JU1200 | C. Braendle[70] |
| NIC664 | *C. elegans* | | F2 RIL(64) JU751 x JU1200 | C. Braendle[70] |
| NIC665 | *C. elegans* | | F2 RIL(65) JU751 x JU1200 | C. Braendle[70] |
| NIC666 | *C. elegans* | | F2 RIL(66) JU751 x JU1200 | C. Braendle[70] |
| NIC667 | *C. elegans* | | F2 RIL(67) JU751 x JU1200 | C. Braendle[70] |
| NIC668 | *C. elegans* | | F2 RIL(68) JU751 x JU1200 | C. Braendle[70] |
| NIC671 | *C. elegans* | | F2 RIL(71) JU751 x JU1200 | C. Braendle[70] |
| NIC675 | *C. elegans* | | F2 RIL(75) JU751 x JU1200 | C. Braendle[70] |
| NIC678 | *C. elegans* | | F2 RIL(78) JU751 x JU1200 | C. Braendle[70] |
| NIC680 | *C. elegans* | | F2 RIL(80) JU751 x JU1200 | C. Braendle[70] |
| NIC681 | *C. elegans* | | F2 RIL(81) JU751 x JU1200 | C. Braendle[70] |
| NIC685 | *C. elegans* | | F2 RIL(85) JU751 x JU1200 | C. Braendle[70] |
| NIC686 | *C. elegans* | | F2 RIL(86) JU751 x JU1200 | C. Braendle[70] |
| NIC687 | *C. elegans* | | F2 RIL(87) JU751 x JU1200 | C. Braendle[70] |

| | | | | |
|---|---|---|---|---|
| NIC691 | *C. elegans* | | F2 RIL(91) JU751 x JU1200 | C. Braendle[70] |
| NIC692 | *C. elegans* | | F2 RIL(92) JU751 x JU1200 | C. Braendle[70] |
| NIC694 | *C. elegans* | | F2 RIL(94) JU751 x JU1200 | C. Braendle[70] |
| NIC696, | *C. elegans* | | F2 RIL(96) JU751 x JU1200 | C. Braendle[70] |
| NIC697 | *C. elegans* | | F2 RIL(97) JU751 x JU1200 | C. Braendle[70] |
| NIC698 | *C. elegans* | | F2 RIL(98) JU751 x JU1200 | C. Braendle[70] |
| NIC699 | *C. elegans* | | F2 RIL(99) JU751 x JU1200 | C. Braendle[70] |
| NIC700 | *C. elegans* | | F2 RIL(100) JU751 x JU1200 | C. Braendle[70] |
| NIC702 | *C. elegans* | | F2 RIL(102) JU751 x JU1200 | C. Braendle[70] |
| NIC703 | *C. elegans* | | F2 RIL(103) JU751 x JU1200 | C. Braendle[70] |
| NIC704 | *C. elegans* | | F2 RIL(104) JU751 x JU1200 | C. Braendle[70] |
| NIC705 | *C. elegans* | | F2 RIL(105) JU751 x JU1200 | C. Braendle[70] |
| NIC707 | *C. elegans* | | F2 RIL(107) JU751 x JU1200 | C. Braendle[70] |
| NIC709 | *C. elegans* | | F2 RIL(109) JU751 x JU1200 | C. Braendle[70] |
| NIC710 | *C. elegans* | | F2 RIL(110) JU751 x JU1200 | C. Braendle[70] |
| NIC711 | *C. elegans* | | F2 RIL(111) JU751 x JU1200 | C. Braendle[70] |
| NIC713 | *C. elegans* | | F2 RIL(113) JU751 x JU1200 | C. Braendle[70] |
| NIC716 | *C. elegans* | | F2 RIL(116) JU751 x JU1200 | C. Braendle[70] |
| NIC717 | *C. elegans* | | F2 RIL(117) JU751 x JU1200 | C. Braendle[70] |
| NIC721 | *C. elegans* | | F2 RIL(121) JU751 x JU1200 | C. Braendle[70] |
| NIC727 | *C. elegans* | | F2 RIL(127) JU751 x JU1200 | C. Braendle[70] |
| NIC728 | *C. elegans* | | F2 RIL(128) JU751 x JU1200 | C. Braendle[70] |
| NIC729 | *C. elegans* | | F2 RIL(129) JU751 x JU1200 | C. Braendle[70] |
| NIC730 | *C. elegans* | | F2 RIL(130) JU751 x JU1200 | C. Braendle[70] |
| NIC731 | *C. elegans* | | F2 RIL(131) JU751 x JU1200 | C. Braendle[70] |
| NIC734 | *C. elegans* | | F2 RIL(134) JU751 x JU1200 | C. Braendle[70] |
| NIC735 | *C. elegans* | | F2 RIL(135) JU751 x JU1200 | C. Braendle[70] |
| NIC736 | *C. elegans* | | F2 RIL(136) JU751 x JU1200 | C. Braendle[70] |
| NIC739 | *C. elegans* | | F2 RIL(139) JU751 x JU1200 | C. Braendle[70] |
| NIC740 | *C. elegans* | | F2 RIL(140) JU751 x JU1200 | C. Braendle[70] |
| NIC741 | *C. elegans* | | F2 RIL(141) JU751 x JU1200 | C. Braendle[70] |
| NIC742 | *C. elegans* | | F2 RIL(142) JU751 x JU1200 | C. Braendle[70] |
| NIC1671 | *C. elegans* | *cgbIR1018* | Near isogenic line *(II:8429315-12658941, JU1200>JU751)* | This study |
| NIC1672 | *C. elegans* | *cgbIR1019* | Near isogenic line *(II: 4185688-11477934, JU1200>JU751)* | This study |
| NIC1673 | *C. elegans* | *cgbIR1020* | Near isogenic line *(II: 5754192-9802124, JU1200>JU751)* | This study |

| | | | | |
|---|---|---|---|---|
| NIC1675 | *C. elegans* | *cgbIR1022* | Near isogenic line *(II: 7239344-11736927, JU1200>JU751)* | This study |
| NIC1676 | *C. elegans* | *cgbIR1023* | Near isogenic line *(II: 11477934-11736927, JU1200>JU751)* | This study |
| NIC1697 | *C. elegans* | *cgbIR1027* | Near isogenic line *(II: 4783409-9867926, JU751>JU1200)* | This study |
| NIC1701 | *C. elegans* | *cgbIR1028* | Near isogenic line *(II: 5968491-11736927, JU751>JU1200)* | This study |
| NIC1702 | *C. elegans* | *cgbIR1029* | Near isogenic line *(II: 10076380-11736927, JU751>JU1200)* | This study |
| NIC1713 | *C. elegans* | *cgbIR1028;cgb1007* | CRISPR line (*cgb1007>NIC1701)* | This study |
| NIC1714 | *C. elegans* | *cgbIR1028;cgb1007* | CRISPR line (*cgb1007>NIC1701)* | This study |
| NIC1715 | *C. elegans* | *cgbIR1028;cgb1007* | CRISPR line (*cgb1007>NIC1701)* | This study |
| NIC1716 | *C. elegans* | *cgbIR1019;cgb1008* | CRISPR line (*cgb1008>NIC1672)* | This study |
| NIC1717 | *C. elegans* | *cgbIR1019;cgb1008* | CRISPR line (*cgb1008>NIC1672)* | This study |
| NIC1719 | *C. elegans* | *cgbIR1019;cgb1008* | CRISPR line (*cgb1008>NIC1672)* | This study |
| NIC1720 | *C. elegans* | *cgb1007 (in JU1200)* | NIC1713 backcrossed to JU1200 to remove *cgb1008* | This study |
| NIC1723 | *C. elegans* | *cgb1008 (in JU751)* | NIC1716 backcrossed to JU751 to remove *cgb1007* | This study |
| NIC1724 | *C. elegans* | *cgb1008 (in JU751)* | NIC1717 backcrossed to JU751 to remove *cgb1007* | This study |
| NIC1725 | *C. elegans* | *cgb1008 (in JU751)* | NIC1718 backcrossed to JU751 to remove *cgb1007* | This study |
| NIC2091 | *C. elegans* | *cgbIR1034* | Near isogenic line *(QII, JU751>JU1200)* | This study |
| MY16 | *C. elegans* | | wild isolate | *C. elegans* Natural Diversity Resource (https://elegansvariation.org) |
